# Over-Representation of Potential SP4 Target Genes within Schizophrenia-Risk Genes

**DOI:** 10.1101/2021.07.14.452377

**Authors:** Xianjin Zhou

## Abstract

Reduction of Sp4 expression causes age-dependent hippocampal vacuolization and many other intermediate phenotypes of schizophrenia in Sp4 hypomorphic mice. Recent human genetic studies from both the Schizophrenia Exome Sequencing Meta-Analysis (SCHEMA) and the Genome-Wide Association Study (GWAS) validated SP4 as a schizophrenia-risk gene over the exome-wide or the genome-wide significance. Truncation of human SP4 gene has an odds ratio of 9.37 (3.38-29.7) for schizophrenia. Despite successful identification of many schizophrenia-risk genes, it is unknown whether and how these risk genes may interact with each other in the development of schizophrenia. By taking advantage of the specific localization of the GC-boxes bound by SP4 transcription factors, I analyzed the relative abundance of these GC-boxes in the proximal promoter regions of schizophrenia-risk genes. I found that the GC-box containing genes are significantly over-represented within schizophrenia-risk genes, suggesting that SP4 is not only a high-risk gene for schizophrenia, but may also act as a hub of network in regulation of many other schizophrenia-risk genes via these GC-boxes in the pathogenesis of schizophrenia.

## Introduction

Sp4, a homologue of the *Drosophila buttonhead* (*btd*) gene, belongs to the Sp1 family of transcription factors (Supp et al., 1996). Sp1, an ubiquitous transcription factor, recognizes the DNA sequence 5’-GGGCGG-3’ termed GC-box that is often found ~40-100 nucleotides upstream of the transcription start site (TSS) of a variety of genes, and is critical for activation of their expression (Kadonaga, 1986). Both Sp1 and Sp4 bind the same GC-box with the same affinity (Hagen et al., 1992). In brain, however, Sp4 and Sp1 are specifically expressed in neuronal and glial cells, respectively (Mao et al., 2007; Zhou et al., 2005). We found that reduction of Sp4 expression in Sp4 hypomorphic mice causes age-dependent hippocampal vacuolization, reduced hippocampal LTP, and many behavioral abnormalities related to schizophrenia such as deficits in prepulse inhibition, deficient learning and memory, hypersensitivity to NMDAR antagonists, etc (Ji et al., 2013; Wang et al., 2018; Zhou et al., 2005; Zhou et al., 2010). A complete absence of Sp4 in Sp4 knockout mice impairs postnatal development of hippocampal dentate gyrus with a reduced number of dentate granule cells and a smaller dentate size (Zhou et al., 2007). We and Tam et al first reported that human SP4 gene is sporadically deleted in schizophrenia patients (Tam et al., 2010; Zhou et al., 2010), and its SNPs associate with both bipolar disorder and schizophrenia (Zhou et al., 2009).

In 2020, the largest GWAS study of 69,369 individuals with schizophrenia and 236,642 controls identified common variant associations at 270 distinct loci (Schizophrenia Working Group of the Psychiatric Genomics Consortium, 2020). Many genes were prioritized after fine mapping of the loci. These genes are predominantly expressed in neurons, but not in glia, and regulate synaptic functions. On the other hand, The Schizophrenia Exome Sequencing Meta-Analysis (SCHEMA) Consortium, the largest exome sequencing project with 24,248 cases and 97,322 controls, identified 10 truncated genes above the exome-wide significance that confer high risks for schizophrenia (odds ratios 3 - 50) (Tarjinder Singh, 2020). Among these top 10 risk genes, truncation of the SP4 gene has an odds ratio of 9.37 (3.38-29.7) for schizophrenia. Importantly, SP4 and GRIN2A (encoding NMDAR2A subunit) are the only two genes shared between the top 10 risk genes from the SCHEMA and the GWAS prioritized genes. This not only enhances the confidence of the two risk genes; but also implicates that more subtle common variants of these two genes with a partial loss-of-function are likely involved in the pathogenesis of schizophrenia in the general population of patients. A key question however that remains to be investigated is whether and how these risk genes may interact with each other in the development of schizophrenia.

Out of the top 10 high-risk genes from the SCHEMA, Sp4 and RB1CC1 are the only two DNA-binding transcription factors. Interestingly, RB1CC1 binds the same GC-box sequence GGGCGG but more upstream of TSS than Sp4 (Ikebuchi et al., 2009), highlighting importance of the GC-box in the regulation of genes involved in the pathogenesis of schizophrenia. In brain, Sp4 and a majority of other schizophrenia-risk genes are specifically expressed in neurons, but not glia. Does Sp4 control expression of many of these schizophrenia-risk genes via the GC-boxes in their proximal promoter regions in neuronal cells, given that the GC-box can be readily found in the 5’-regulatory regions of a variety of genes? Here, I examined the relative abundance of the GC-boxes in the promoters of susceptibility genes for schizophrenia, autism spectrum disorders (ASD), and developmental disorder/intellectual disability (DD/ID).

## Materials and Methods

### Eukaryotic promoter database

Eukaryotic promoter database (EPD) (https://epd.epfl.ch//index.php) has a collection of 16,455 genes across human genome (hg38). The GC-box GGGCGG, the GA-box GGGAGG, the MAZ-binding motif CCCCTCC, and the KLF13-binding motif ACGCCC, were searched on both DNA strands in gene promoters using function of either Motif Enrichment or Motif Discovery in EPD. The numbers of the GC-boxes were counted for each gene. To calculate the GC content of the promoter region, either G or C was counted on both DNA strands using Motif Discovery. The numbers of the GC-boxes and the GC content were analyzed using R programming.

### Susceptibility genes for schizophrenia, ASD, and DD/ID

High-risk genes for schizophrenia were obtained from the SCHEMA (Tarjinder Singh, 2020). Common risk genes for schizophrenia came from the GWAS (Schizophrenia Working Group of the Psychiatric Genomics Consortium, 2020). High-risk genes for ASD were obtained from large scale exome sequencing studies (Satterstrom et al., 2020). Risk genes for DD/ID came from exome sequencing studies (Joanna Kaplanis, 2019; Tarjinder Singh, 2020).

### Statistical analysis

R programming was used for all statistical analyses and correction of multiple comparisons by FDR. Statistical analysis of over-representation of the GC-box containing genes was conducted with cumulative hypergeometric probability tests. Permutation tests (N=9,999) were conducted for statistical analyses of the numbers of the GC-boxes in the proximal promoter regions between different groups of genes. ANOVA or Welch’s *t*-test was used for statistical analysis of the GC-content in the proximal promoter regions between different groups of genes.

## Results

Eukaryotic Promoter Database (EPD) (https://epd.epfl.ch//index.php) hosts 16,455 protein-coding genes across human genome (hg38) with well-characterized TSS. To avoid distortion by different numbers of alternative promoters between genes, the most representative promoter was selected from each gene for analysis. Consistent with Kadonaga’s findings (Kadonaga, 1986), the GC-box is predominantly localized starting from 40 nucleotides upstream of TSS after scanning the promoter region (−500,100) of all 16,455 genes (Figure 1). The sudden drop of the GC-box percentages at 40 nucleotides upstream of TSS separates the proximal promoter from the core promoter that is occupied by RNA polymerase II and associated basal transcription machinery. The 100 nucleotide (−140,−41, red line) window for the GC-box is similar to the (−130,−30) window in previous studies (Yokoyama and Pollock, 2012) and slightly larger than Kadonaga’s (−100,−40) window in early studies, likely due to more accurate TSS mapping of many thousands of genes in the most recent EPD database. In this proximal promoter region, the GC-boxes function as critical Sp1 and Sp4 binding sites for activation of gene expression (Dynan and Tjian, 1983; Hagen et al., 1995). The proximal promoter region (red line) from −140 to −41 was therefore selected as a critical window for analysis of potentially functional GC-boxes in schizophrenia-risk genes, ASD-risk genes, and DD/ID-risk genes.

**Figure 1.**
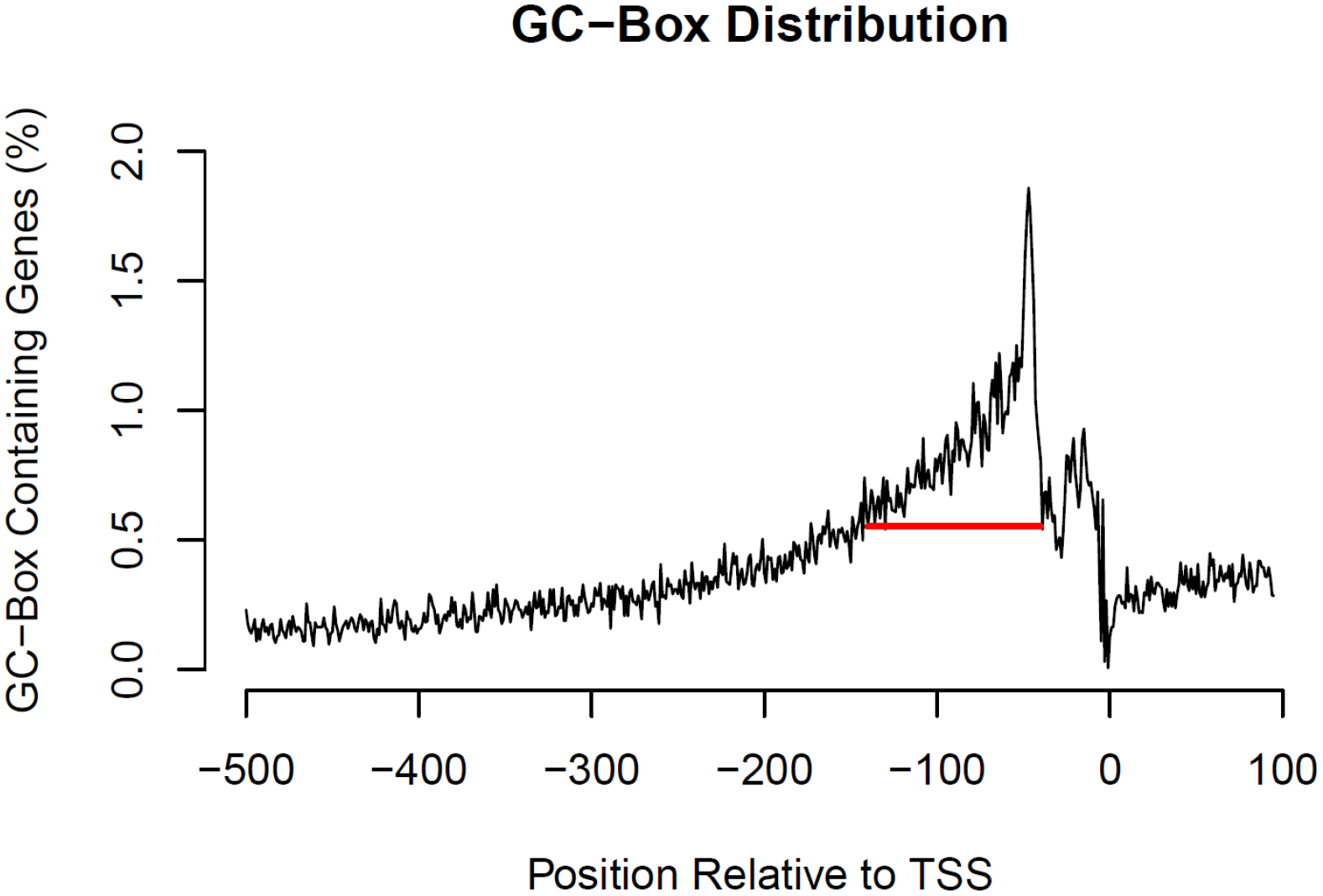
A critical window for the GC-box in regulation of gene expression. The GC-box GGGCGG was bidirectionally scanned across the promoter region (−500,100) of 16,455 human genes in the EPD database using Motif Enrichment with a window size of 6 nucleotides and 1 nucleotide window shift. The relative abundance of the GC-boxes was plotted across the whole region. A sudden drop of the GC-boxes about 40 nucleotides upstream of TSS indicates the boundary between the proximal and the core promoter. A 100 nucleotide (−140,−41, red line) window was selected to ensure a high density of potentially functional GC-boxes.

### Over-representation of GC-box containing genes

After identifying the GC-boxes in each of the 16,455 genes in the EPD, I found that 7,653 genes, ~46.51% of all genes, have at least one GC-box in their proximal promoter (−140,−41) regions (Table 1). This percentage of the GC-box containing genes across human genome is the control baseline for determination of over-representation of the GC-box containing genes within susceptibility genes for schizophrenia, ASD, and DD/ID. Since the top 10 high-risk genes with the exome-wide significance from the SCHEMA are too small for statistical analysis, I expanded the gene list to the top 32 high-risk genes with p value < 10^−4^ (Tarjinder Singh, 2020), presumably most of the expanded genes will reach the exome-wide significance after sequencing more cases and controls. Not all human genes have been finely mapped with TSS and collected in the EPD database. After searching the 32 high-risk genes in the EPD, 28 genes were matched. Out of the 28 EPD-matched genes, 19 genes, ~67.86% of the matched high-risk genes, have at least one GC-box in their proximal promoter regions. Cumulative hypergeometric probability test suggested that the 19 GC-box containing genes are significantly (p=0.0186) over-represented within the 28 EPD-matched high-risk genes from the SCHEMA than the 7,653 GC-box containing genes within all 16,455 human genes. If the gene list is further expanded to the top 61 high-risk genes with p value < 10^−3^ from the SCHEMA, 47 out of the 61 genes are matched in the EPD. 31 out of the 47 EPD-matched genes, ~65.96% of all these genes, contain at least one GC-box in their proximal promoter regions. Again, the 31 GC-box containing genes are significantly (p=0.0056) over-represented within the 47 EPD-matched high-risk genes from the SCHEMA than the 7,653 GC-box containing genes within all 16,455 human genes. In contrast to the rare high-risk genes identified by the SCHEMA, the GWAS found hundreds of common low-risk genes for schizophrenia with the genome-wide significance (Schizophrenia Working Group of the Psychiatric Genomics Consortium, 2020). After fine-mapping of the loci, 69 protein-coding genes were prioritized. 63 out of the 69 genes are matched in the EPD. 36 out of the 63 EPD-matched genes have at least one GC-box in their proximal promoter regions. Cumulative hypergeometric probability test suggested a trend (p=0.058) of over-representation of the 36 GC-box containing genes within the 63 EPD-matched genes prioritized by the GWAS. If all 643 GWAS prioritized genes were used for the same analysis, the number of GC-box containing genes is also significantly over-represented (p=0.0306). In conclusion, the GC-box containing genes are significantly over-represented within the schizophrenia-risk genes identified from both the SCHEMA and the GWAS studies. Are the GC-box containing genes also over-represented within high-risk genes for other psychiatric disorders? I analyzed the proportion of the GC-box containing genes within the high-risk genes for autistic spectrum disorders (ASD) from large-scale exome sequencing (Satterstrom et al., 2020). There is no over-representation of the GC-box containing genes (p=0.45), suggesting that the GC-boxes in the proximal promoter regions (−140,−41) are functionally less dominant in the regulation of the ASD-risk genes. Risk genes for developmental disorder/intellectual disability (DD/ID) (Joanna Kaplanis, 2019; Tarjinder Singh, 2020) were also analyzed. 141 GC-box containing genes are significantly (p=0.005) over-represented within the 258 matched high-risk genes for DD/ID than the 7,653 GC-box containing genes within all 16,455 human genes. After correcting multiple comparisons with FDR, significant over-representation of the GC-box containing genes remains in both schizophrenia and DD/ID.

**Table 1.** GC-Boxes in Proximal Promoter Regions of Risk Genes for SCZ, ASD, DD/ID.

|  | **Total**<br>**Genes** | **EPD**<br>**Matched**<br>**Genes** | **GC-Box**<br>**Containing**<br>**Genes** | **GC-Box**<br>**Gene %** | **Cumulative**<br>**Hypergeometric**<br>**Probability (p)** | **FDR-**<br>**adjusted**<br>**p Value** |
| --- | --- | --- | --- | --- | --- | --- |
| **EPD All Genes (hg38)** | 16455 | 16455 | 7653 | 46.51 |  |  |
| **SCZ Genes (SCHEMA)** |  |  |  |  |  |  |
| p value < 1E-4 | 32 | 28 | 19 | **67.86** | **0.0186\*** | **0.037\*** |
| p value < 1E-3 | 61 | 47 | 31 | **65.96** | **0.0056\*\*** | **0.017\*** |
| **SCZ Genes (GWAS)** |  |  |  |  |  |  |
| **FINEMAP Prioritized Genes** | 69 | 63 | 36 | **57.14** | **0.058** | **0.07** |
| **All Prioritized Genes** | 643 | 357 | 184 | **51.54** | **0.0306\*** | **0.046\*** |
| **ASD Genes** | 102 | 88 | 42 | 47.73 | 0.4503 | 0.45 |
| **DD/ID Genes** | 299 | 258 | 141 | **54.65** | **0.005\*\*** | **0.017\*** |

The GC-box evolves to be the dominant motif bound by Sp1 and Sp4 transcription factors with the highest affinity in eutherian mammals including mouse and human, whereas the ancestral GA-box GGGAGG is more common for Sp1 and Sp4 transcription factors in other vertebrates such as fish and marsupials (Yokoyama and Pollock, 2012). The GC-box has 3 times higher affinity for Sp1/Sp4 transcription factors than the GA-box (Letovsky and Dynan, 1989). Proportion of the GA-box containing genes within the risk genes for schizophrenia, ASD, and DD/ID was analyzed with exactly the same method (Table 2). There is no over-representation of the GA-box containing genes within the schizophrenia-risk genes or the ASD-risk genes except for the DD/ID-risk genes. Since the GA-box is almost identical to the binding motif (CCCCTCC) of the transcription factor MAZ, the same pattern of the overrepresentation of MAZ motif-containing genes was observed in the DD/ID-risk genes (Supplemental Table S1). It is possible that the two over-represented similar motifs may be mainly selected by MAZ. As expected, the binding motif (ACGCCC) of transcription factor KLF13 does not show enrichment in any group of the risk genes (Supplemental Table S2), supporting that over-representation of the GC-box containing genes within the schizophrenia-risk genes or DD/ID-risk genes is not a random effect.

**Table 2.** GA-Boxes in Proximal Promoter Regions of Risk Genes for SCZ, ASD, DD/ID.

|  | Total<br>Genes | EPD<br>Matched<br>Genes | GA-Box<br>Containing<br>Genes | GA-Box<br>Gene % | Cumulative<br>Hypergeometric<br>Probability (p) | FDR-adjusted p<br>Value |
| --- | --- | --- | --- | --- | --- | --- |
| **EPD All Genes (hg38)** | 16455 | 16455 | 5485 | 33.33 |  |  |
| **SCZ Genes (SCHEMA)** |  |  |  |  |  |  |
| p value < 1E-4 | 32 | 28 | 10 | 35.71 | 0.4645 | 0.4645 |
| p value < 1E-3 | 61 | 47 | 18 | 38.29 | 0.2813 | 0.3375 |
| **SCZ Genes (GWAS)** |  |  |  |  |  |  |
| FINEMAP Prioritized<br>Genes | 69 | 63 | 25 | 39.68 | 0.1738 | 0.3375 |
| All Prioritized Genes | 643 | 357 | 126 | 35.29 | 0.2295 | 0.3375 |
| **ASD Genes** | 102 | 88 | 37 | 42.05 | 0.0539 | 0.1617 |
| **DD/ID Genes** | 299 | 258 | 118 | **45.74** | **0.00002\*\*\*** | **0.0001\*\*\*** |

### Numbers of the GC-boxes in the GC-box containing genes

Since the GC-box containing genes are over-represented within schizophrenia-risk genes and DD/ID-risk genes, I examined whether the GC-boxes may also be enriched in the GC-box containing risk genes. The GC-boxes were identified and counted in the proximal promoter regions (−140,−41) of the 7,653 GC-box containing genes in the EPD (Supplemental Table S3). Schizophrenia-risk genes from the SCHEMA and the GWAS, ASD-risk genes, and DD/ID-risk genes were extracted (Supplemental Table S3). Distribution of the numbers of the GC-boxes in the GC-box containing genes was boxplotted (Figure 2A). Due to non-normal distribution, permutation test (N=9,999) was conducted for statistical analysis (Supplemental Table S4). No significant differences were found between the control EPD group and each group of disease-risk genes, suggesting that there is no enrichment of the GC-boxes in the proximal promoter regions of the GC-box containing risk-genes for schizophrenia, ASD, and DD/ID

**Figure 2.**
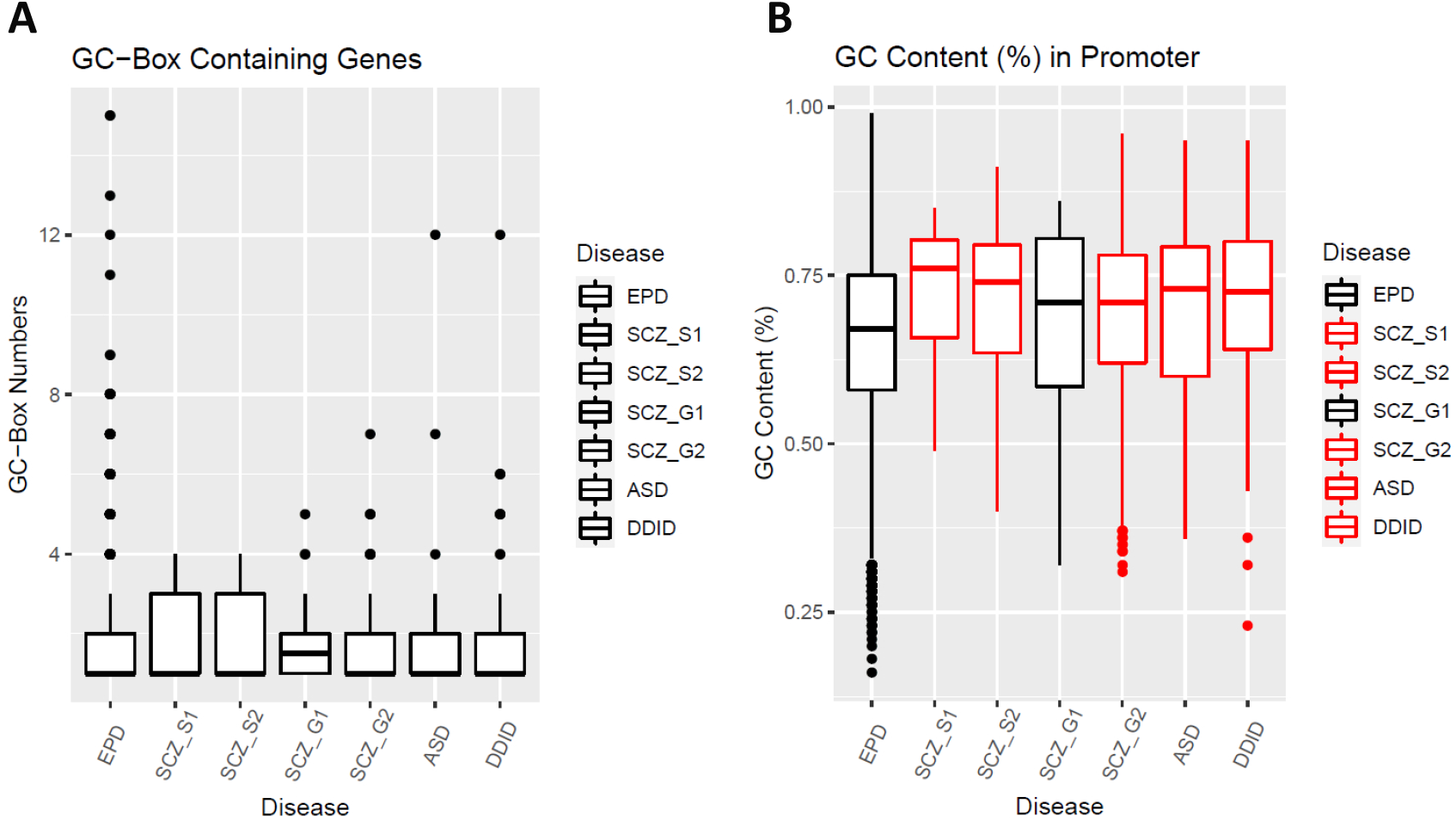
The GC-boxes and the GC-content in the proximal promoter region (−140,−41). SCZ_S1 (SCHEMA1, risk-genes with p < 10^−4^), SCZ_S2 (SCHEMA2, risk-genes with p < 10^−3^), SCZ_G1 (GWAS1, FINEMAP prioritized genes), SCZ_G2 (GWAS2, all prioritized genes). **(A)** The numbers of the GC-boxes in the proximal promoter region (−140,−41) of the GC-box containing genes was boxplotted in different groups of genes. Permutation tests showed no significant difference between the EPD group and any of the disease groups. **(B)** The GC content in the proximal promoter region (−140,−41) was calculated for each gene. Welch’s *t*-tests were used for statistical analysis (Supplemental Table S4). All of the risk-gene groups except SCZ_G1 (black) have a significantly higher GC content than the EPD group after FDR correction of multiple comparisons.

### Association of the GC-box with the GC content

High GC content may generate the GC-box by chance in the promoter. Does the over-representation of the GC-box containing genes result from higher GC content in the proximal promoter regions of schizophrenia-risk genes and DD/ID-risk genes? The GC content (%) in the proximal promoter region (−140,−41) was calculated for all 16,455 genes including the 7,653 GC-box containing genes in the EPD (Supplemental Table S3). Schizophrenia-risk genes from the SCHEMA and the GWAS, ASD-risk genes, and DD/ID-risk genes were extracted (Supplemental Table S3). The GC-content in the proximal promoter region (−140,−41) was boxplotted (Figure 2B). All groups of genes except SCZ_G1 have a significantly higher GC content than the control EPD genes by Welch’s t-tests and FDR correction (Supplemental Table S4). A high GC content of the ASD-risk genes does not produce more GC-boxes in their proximal promoter regions (Table I); suggesting that over-representation of the GC-box containing genes in the schizophrenia-risk genes cannot be simply attributed to the increased GC content. If the GC content was separately boxplotted in the GC-box containing genes or non-GC-box genes, the GC-box containing genes have a significantly higher GC content than non-GC-box genes across all groups regardless of diseases (Supplemental Figure S1). All together, the presence of the GC-boxes will increase the GC content, but a high GC content does not necessarily produce more GC-boxes. Over-representation of the GC-box containing genes likely contributes most to the higher GC content in the proximal promoter regions of the schizophrenia-risk genes and the DD/ID-risk genes.

### The GC-boxes more upstream of TSS

In contrast to Sp1/Sp4 transcription factors that recognize the GC-boxes in the proximal promoter region (−140,−41), the GC-boxes more upstream of this region can be bound by other transcription factors such as RB1CC1, another high-risk gene for schizophrenia from the SCHEMA. These transcription factors are however less well studied; and their binding GC-boxes scatter across a larger area of the 5’-regulatory region. Such more flexible localizations in a large region make analysis more difficult and arbitrary. To get a glimpse of potential involvement of these more upstream GC-boxes in schizophrenia, I plotted the percentages of the GC-boxes in the promoters of susceptibility genes for schizophrenia, ASD, and DD/ID (Figure 3). In addition to the major peak of the GC-boxes in the proximal promoter region (−140,−41) across all schizophrenia-risk genes, there appears a second peak of more upstream GC-boxes (−340,−240) in the high-risk genes from SCHEMA2. This second GC-box peak is however absent in schizophrenia-risk genes prioritized by the GWAS. In high-risk susceptibility genes for ASD and DD/ID, the frequency of these more upstream GC-boxes is higher than that of all 16,455 genes across a large 5’-regulatory region. Significance of these upstream GC-boxes remains to be investigated in the regulation of expression of these genes.

**Figure 3.**
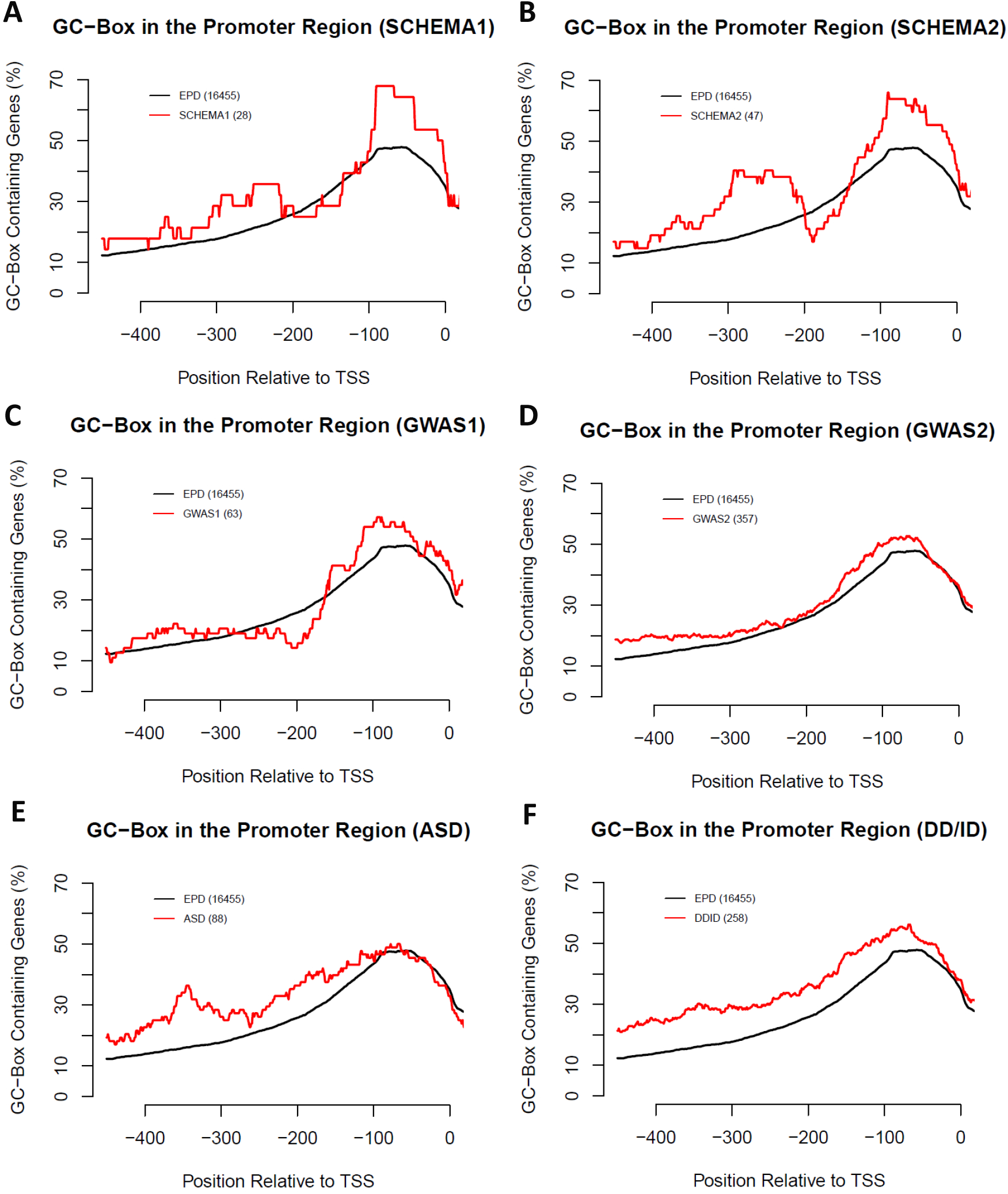
The GC-boxes upstream of the proximal promoter region (−140,−41). Due to small sample sizes of schizophrenia-risk genes, a large window size (100 nucleotides) was used to scan the frequency of the GC-boxes across the promoter regions (−500,50) with 1 nucleotide window shift. The percentages of the GC-box containing genes in each window were plotted for the control EPD and each group of disease genes in SCHEMA1 **(A)**, SCHEMA2 **(B)**, GWAS1 **(C)**, GWAS2 **(D)**, ASD **(E)**, and DD/ID **(F)**.

## Discussion

The SCHEMA identified 10 high-risk genes for schizophrenia above the exome-wide significance. Out of the top 10 high-risk genes, Sp4 and RB1CC1, the only two DNA binding transcription factors, bind the same GC-box in regulation of genes involved in the pathogenesis of schizophrenia. The GC-box containing genes are significantly over-represented in the susceptibility genes for schizophrenia from both the SCHEMA and the GWAS studies, suggesting that Sp4 is not only a high-risk gene for schizophrenia, but may also act as a hub of network in the regulation of many other schizophrenia-risk genes via these GC-boxes in neuronal cells. Such a Sp4-controlled network of different schizophrenia-risk genes may contribute to the over-representation of the GC-box containing genes. In the GC-box containing risk-genes for schizophrenia or DD/ID, the numbers of the GC-boxes are not significantly increased in their proximal promoter regions, indicating a lack of synergistic effects from multiple GC-boxes in activation of gene expression. Interestingly, Sp4 distinguishes itself from Sp1 by lack of such synergistic effects from multiple adjacent GC-boxes (Hagen et al., 1995).

During evolution of eutherian mammals, the GC-box rises quickly with co-evolved Sp1 and Sp4 variants after split from marsupials (Yokoyama and Pollock, 2012). The ancestral GA-boxes for Sp1/Sp4 transcription factors decrease by genetic drift. In human, there are still a significant large number of the GA-boxes in the proximal promoter regions (−140,−41). Unlike the GC-boxes, however, there is no over-representation of the GA-box containing genes within the schizophrenia-risk genes likely due to lack of selective pressure for the GA-boxes by Sp4 and other possible transcription factors. Surprisingly, over-representation of the GA-box containing genes was still observed within the DD/ID-risk genes. Given that the MAZ transcription factor also binds the GA-box that is similar to the MAZ-binding motif, it is plausible that MAZ may regulate genes involved in the development of DD/ID and therefore causes the over-representation of both the GA-boxes and the MAZ-binding motifs. It cannot be ruled out, however, that the GA-boxes, despite with a lower affinity than the GC-boxes, may still be used in a limited capacity for Sp1/Sp4/Sp3 transcription factors depending on gene and cellular context in the development and diseases of the central nervous system.

Not all GC-boxes in the proximal promoter region will be bound by Sp4 transcription factors in neuronal cells. Whether a specific GC-box in a schizophrenia-risk gene is indeed occupied by Sp4 has to be determined by experiments such as chromatin immunoprecipitation and sequencing (ChIP-seq). It should be kept in mind that other Sp1 family of transcription factors such as Sp3 may also recognize the GC-boxes in the proximal promoter regions to activate or repress gene expression.

Over-representation of the GC-box containing genes appears more prominent within the high-risk genes from the SCHEMA than within the common low-risk genes from the GWAS. A possible explanation is that the schizophrenia-risk genes from the GWAS carry much smaller effects and are less certain, particularly for the 643 genes (GWAS2) prioritized by various methodologies and assumptions. Over-representation of the GC-box containing genes is also observed within the risk-genes for DD/ID, but not ASD. Given that mouse Sp4 mutation impairs hippocampal development and causes severe deficits in learning and memory, it is not surprising that the GC-boxes are over-represented in the proximal promoter regions of the risk genes involved in the development of DD/ID.

The frequency of the GC-boxes peaks at the proximal promoter region (−140,−41) where Sp4 binds to activate gene expression in schizophrenia-risk genes. A second peak of more upstream GC-boxes is suggested in the high-risk genes from the SCHEMA. Given that RB1CC1, a high risk gene for schizophrenia, binds the more upstream GC-boxes, it is possible that these upstream GC-boxes may also be involved in the pathogenesis of schizophrenia. Identification of more high-risk schizophrenia genes will help confirm whether these upstream GC-boxes are also significantly over-represented. In contrast to the schizophrenia-risk genes, susceptibility genes for ASD and DD/ID have more upstream GC-boxes across a large promoter region. It is unclear whether they may have any functional significance or are just produced by chance due to a high GC content in the promoters of these risk genes. However, a high GC content itself could be a consequence of selective pressure by transcription factors recognizing GC-rich motifs in regulation of these risk genes in the pathogenesis of ASD and DD/ID.

