## Supplementary figures and images for "Over-Representation of Potential SP4 Target Genes within Schizophrenia-Risk Genes"

### Supplemental Figure S1

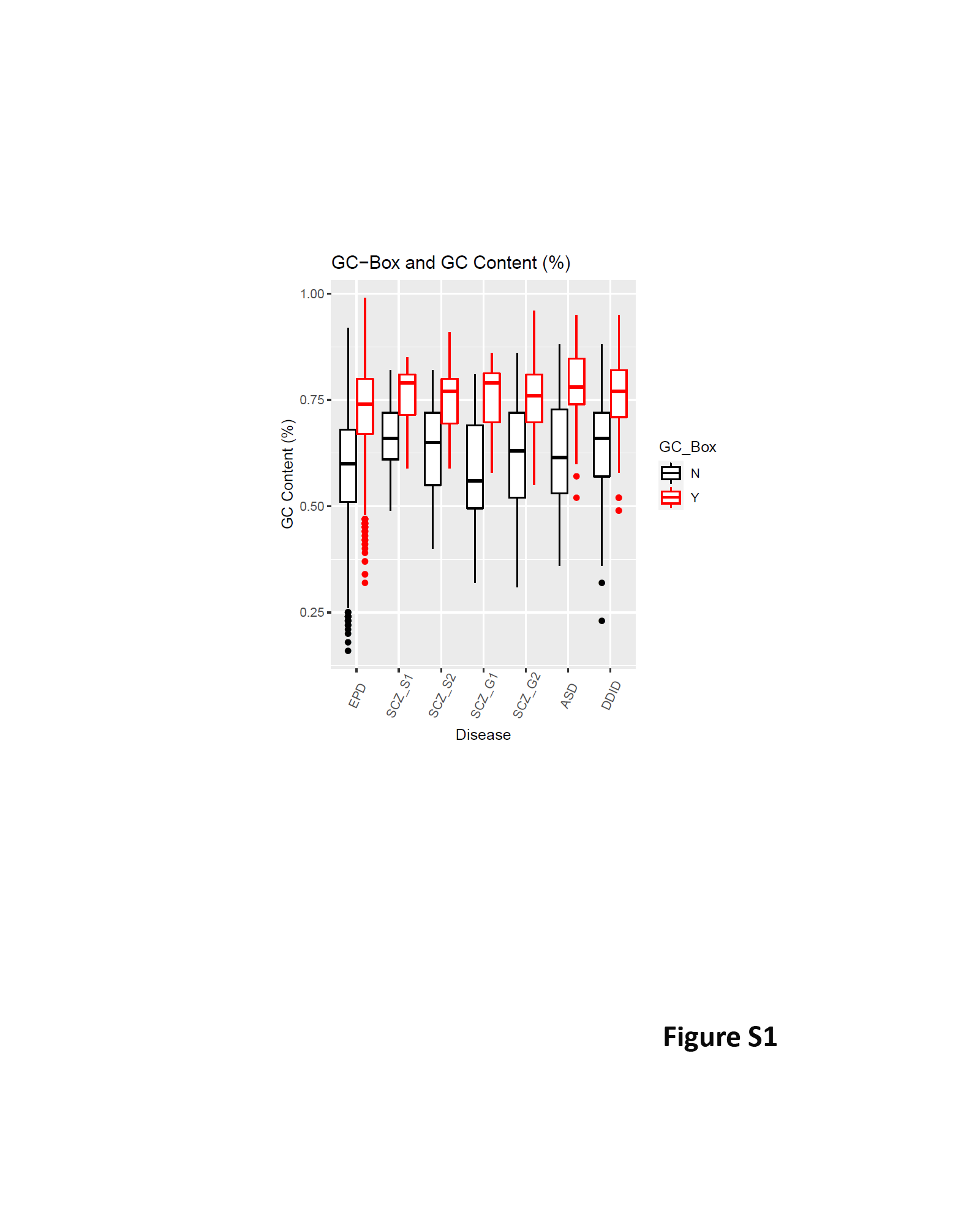
