## Supplemental Materials for "Over-Representation of Potential SP4 Target Genes within Schizophrenia-Risk Genes"

**Supplemental Figure Legend**

**Supplemental Figure S1. The GC-box containing genes have a higher GC content in their proximal promoter regions.** The genes containing at least one GC-box (Y) and the genes without the GC-box (N) were separated for boxplotting their GC contents in different groups of genes. The GC-box containing genes have a significantly higher GC content (F(1,17282)=7739.34, p < 2 x 10^-16^) than genes without GC-box across all groups regardless of diseases. SCZ_S1 (SCHEMA1, risk-genes with p < 10^-4^), SCZ_S2 (SCHEMA2, risk-genes with p < 10^-3^), SCZ_G1 (GWAS1, FINEMAP prioritized genes), SCZ_G2 (GWAS2, all prioritized genes).
