## Supplemental Table S1 for "Over-Representation of Potential SP4 Target Genes within Schizophrenia-Risk Genes"

| **Table S1. MAZ Motif in Proximal Promoter Regions of Risk Genes for SCZ, ASD, DD/ID** | | | | | | | | | | | |
| --- | --- | --- | --- | --- | --- | --- | --- | --- | --- | --- | --- |
|  | | **Total Genes** | | **EPD Matched Genes** | | **MAZ Motif Containing Genes** | | **MAZ Motif Gene %** | | **Cumulative Hypergeometric Probability (p)** | **FDR-adjusted p Value** |
| **EPD All Genes (hg38)** | | 16455 | | 16455 | | 2822 | | 17.14 | |  |  |
| **SCZ Genes (SCHEMA)** | |  | |  | |  | |  | |  |  |
| **p value < 1E-4** | | 32 | | 28 | | 5 | | 17.86 | | 0.5380 | 0.4645 |
| **p value < 1E-3** | | 61 | | 47 | | 11 | | 23.40 | | 0.1706 | 0.3375 |
| **SCZ Genes (GWAS)** | |  | |  | |  | |  | |  |  |
| **FINEMAP Prioritized Genes** | | 69 | | 63 | | 12 | | 19.05 | | 0.3946 | 0.3375 |
| **All Prioritized Genes** | | 643 | | 357 | | 57 | | 15.97 | | 0.3023 | 0.3375 |
| **ASD Genes** | | 102 | | 88 | | 20 | | 22.73 | | 0.1082 | 0.1617 |
| **DD/ID Genes** | | 299 | | 258 | | 70 | | **27.13** | | **0.000036***** | **0.0001***** |
