## Supplemental Table S2 for "Over-Representation of Potential SP4 Target Genes within Schizophrenia-Risk Genes"

| **Table S2. KLF13 Motif in Proximal Promoter Regions of Risk Genes for SCZ, ASD, DD/ID** | | | | | | | | | | | |
| --- | --- | --- | --- | --- | --- | --- | --- | --- | --- | --- | --- |
|  | | **Total Genes** | | **EPD Matched Genes** | | **KLF Motif Containing Genes** | | **KLF Motif Gene %** | | **Cumulative Hypergeometric Probability (p)** | **FDR-adjusted p Value** |
| **EPD All Genes (hg38)** | | 16455 | | 16455 | | 1664 | | 10.11 | |  |  |
| **SCZ Genes (SCHEMA)** | |  | |  | |  | |  | |  |  |
| **p value < 1E-4** | | 32 | | 28 | | 2 | | 7.14 | | 0.4513 | 0.5416 |
| **p value < 1E-3** | | 61 | | 47 | | 6 | | 12.76 | | 0.3381 | 0.5072 |
| **SCZ Genes (GWAS)** | |  | |  | |  | |  | |  |  |
| **FINEMAP Prioritized Genes** | | 69 | | 63 | | 6 | | 9.52 | | 0.5438 | 0.5438 |
| **All Prioritized Genes** | | 643 | | 357 | | 40 | | 11.20 | | 0.2685 | 0.5072 |
| **ASD Genes** | | 102 | | 88 | | 4 | | 4.55 | | **0.0492** | 0.2952 |
| **DD/ID Genes** | | 299 | | 258 | | 22 | | 8.53 | | 0.2311 | 0.5072 |
