## Supplemental Table S4 for "Over-Representation of Potential SP4 Target Genes within Schizophrenia-Risk Genes"

Permutation tests (N=9,999) for the GC box numbers

```
      Disease   p_value
1 SCZ_SCHEMA1 0.6167617
2 SCZ_SCHEMA2 0.8805881
3   SCZ_GWAS1 0.9399940
4   SCZ_GWAS2 0.2154215
5         ASD 0.1905191
6        DDID 0.4580458
```

| **Welch's t-tests for the GC-content and FDR adjusted p values** | | | | | | | |
| --- | --- | --- | --- | --- | --- | --- | --- |
|  | | **SCZ_S1** | **SCZ_S2** | **SCZ_G1** | **SCZ_G2** | **ASD** | **DD/ID** |
| **mean of EPD** | | 0.6563 | 0.6563 | 0.6563 | 0.6563 | 0.6563 | 0.6563 |
| **mean of risk genes** | | 0.7254 | 0.7077 | 0.6771 | 0.6840 | 0.6959 | 0.7059 |
| **ci.lower** | | -0.1068 | -0.0830 | -0.0569 | -0.0416 | -0.0682 | -0.0643 |
| **ci.upper** | | -0.0312 | -0.0196 | 0.0153 | -0.0138 | -0.0109 | -0.0348 |
| **p.value** | | **0.0009** | **0.0021** | 0.2533 | **0.0001** | **0.0073** | **1.98E-10** |
| **p_FDR** | | **0.0017** | **0.0031** | 0.2533 | **0.0003** | **0.0088** | **1.19E-09** |
